# Decreased *in vitro* dihydroartemisinin sensitivity in malaria parasites infecting sickle cell disease patients

**DOI:** 10.1101/2022.04.29.490119

**Authors:** Albert A. Gnondjui, Offianan A. Toure, Beranger A. Ako, Tossea S. Koui, Stanislas E. Assohoun, Eric A. Gbessi, Landry T. N’guessan, Karim Tuo, Sylvain Beourou, Serge-Brice Assi, Francis A. Yapo, Ibrahima Sanogo, Ronan Jambou

## Abstract

**Background:** Partial ACTs treatment failure in *Plasmodium falciparum* malaria has been previously reported in sickle cell patients. The main purpose of this study was to investigate the *in vitro* susceptibility of clinical isolates to DHA to find out hypothesis backing up the reason of this poor therapeutic response.

**Results:** A total of 134 clinical isolates from patients attending health centers in Abidjan with uncomplicated *Plasmodium falciparum* malaria were selected. Hemoglobin HbAS, HbSS, HbAC, HbSC and HbAA were identified. Parasitemia and hemoglobin level at inclusion were lower in sickle cell patients with major forms than in patients with normal phenotype. A significant number of parasites with survival rates ranging from 14.68 to 33.75% were observed in clinical isolates from the SS phenotype. At inclusion, these resistant clinical isolates showed lower parasite densities, and patients had lower red blood cell count and hematocrit levels compared to those with susceptible clinical isolates. A low rate of parasitic growth has more often occurred with AS sickle cell phenotype. However, the decrease in *in vitro* sensitivity to DHA was not associated with Kelch 13-Propeller gene polymorphism.

**Conclusion:** This study highlights an *in vitro* decreased sensitivity to DHA, for clinical isolates collected from sickle cell SS patients living in Abidjan (Côte d’Ivoire), which is not related to the Pfkelch13 gene mutations. These clinical isolates may represent a health threat for sickle cell disease patients especially during crisis. Moreover, these results could suggest additional mechanisms of artemisinin resistance that need to be explored.

## INTRODUCTION

Parasitic infections such as *Plasmodium falciparum* malaria are one of the major causes of morbidity and mortality in patients with major sickle cell disease (SS, SC and CC) (1–5). Indeed, the development of the parasite in sickle cell (6–8), may cause vaso-occlusive crises and increase hemolytic anemia through acute hemolysis episodes (9–12). In endemic areas, malaria is considered to be one of the main causes of frequent hospitalization for sickle cell disease patients (2, 13). Studies showed that sickle cell trait (heterozygous AS) does not prevent against malaria infection but protects against severe malaria (14–16). However, the homozygous status (SS) may be associated with increased susceptibility to malaria (17). Red Blood Cells with abnormal hemoglobin are very different from others in terms of redox potentials and calcium fluxes (18, 19). These ionic changes in red blood cells can induce selection pressure on all proteins involved in the ionic regulation of the parasite (6, 19). These modified intracellular media could also modulate the metabolic pathways involved in artesunate resistance by selecting resistant parasites. This supports development of *in vitro* studies of *P. falciparum* resistance to artemisinin derivatives in sickle cell patients to discriminate between, mutation or transcriptome regulation of the parasite, or partial inactivation of the drug in abnormal red cells.

Artemisinin-based combination therapies (ACTs) resistance emerged in the Mekong region known to have a very high prevalence of hemoglobin E (20–24). Recent studies showed that, *Plasmodium falciparum* resistance to artemisinin and its derivatives at ring stage was based on a quiescence mechanism (25–30). In Asia, this resistance was attributed to mutations in the K13 propeller gene (31–33). However, in Africa more and more reports pointed out a decrease in treatment response not related to mutations in this gene (34–37). In the other side, and despite early description of mutations in the SERCA gene, other genes have been recently described to correlate with the decrease in resistance (38–43). All these genes are involved in the homeostasis of the cell content. Their expression could be highly modulated when facing the very special cytosol content of the abnormal red cells.

In Côte d’Ivoire the prevalence of abnormal hemoglobin is very high (44). Since 2005, Côte d’Ivoire has been using ACTs in the treatment of uncomplicated *P. falciparum* malaria. A very high polymorphism of the K13 propeller gene of parasites obtained from patients with uncomplicated *P*.*falciparum* malaria, was also described (45). Adjei and colleagues in Ghana, as well as Gbessi and colleagues in Côte d’Ivoire reported decreased efficacy of ACT treatments in sickle cell patients correlated with a delay in *Plasmodium falciparum* parasite clearance (37, 46, 47). This suggests a possible resistance of *Plasmodium falciparum* in this population. In the same time Gbessi et al highlighted a larger phenotypic complexity of the parasite populations in patients with SCD than in normal ones. However, the *in vivo* therapeutic efficacy test for ACTs does not allow a direct analysis of parasites response to artemisinin derivatives because the additional effect of patient pre-munition on drug efficacy could mask the detection of chemo-resistant isolates in high transmission area.

The aim of this study was to investigate the *in vitro* susceptibility of parasites to DHA, causing malaria in sickle cell patients. For this purpose, phenotypic tests (Ring Stage Assay and Schizont Maturation Standard Tests) and genotypic test (K13 propeller gene sequence analysis) were carried out to evaluate the susceptibility of these parasites to DHA.

## RESULTS

### Study population

During the study, a total of 1567 patients attended health centers with suspected malaria mild infection. However, most of them (**figure 1**) had either a negative rapid test or a negative thin blood smear. Patients which had already undergone treatment were discarded from the study. A total of 134 patients were enrolled and blood samples were collected. Among them 72 had AA phenotypes, 26 were heterozygotes (AC or AS) and 36 were double mutated (SC or SS) **(figure 1**). Surprisingly no CC was found.

**Figure 1:**
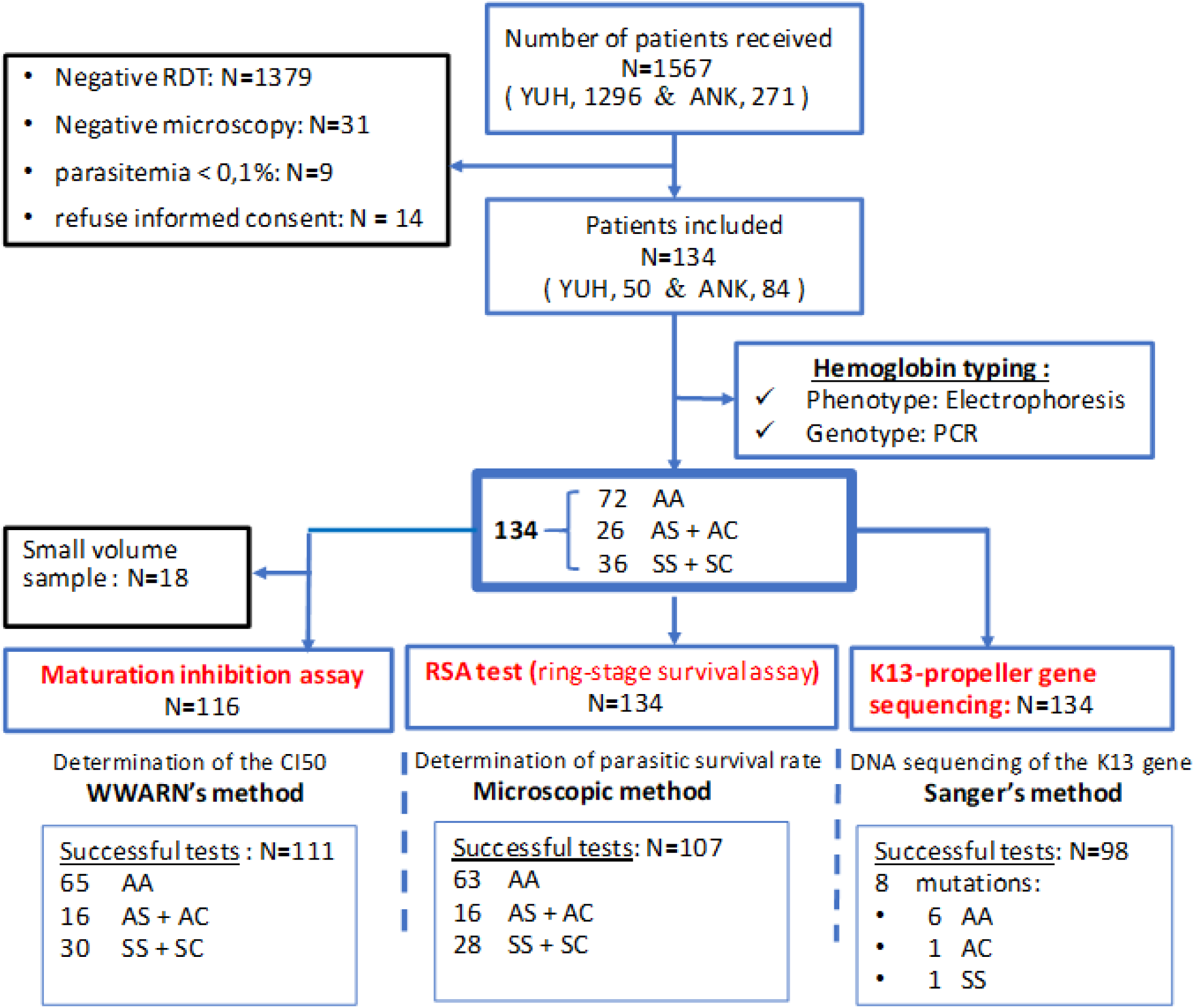
Flow chart of the study.

In the present study, most of the recruited patients were children. Participants with major sickle cell disease harboring vaso-occlusive crisis (CVO) were admitted. They were treated with non-steroidal anti-inflammatory drugs before attending health centers. The mean age in the normal AA phenotype group (12.9 years) was not significantly different from that of the minor AS and AC and major SC and SS (**table 1**). The mean temperature and parasite density at inclusion were lower in patients with HbAS, HbSC and HbSS compared to normal HbAA group (Mann Whitney test, P < 0.001). Likewise, erythrocyte count, hemoglobin and hematocrit were lower at inclusion for HbAS, HbSC and HbSS patients compared to HbAA (Mann Whitney test, P < 0.001). A difference was also found between groups like HbAC and HbSC (Mann Whitney test, P <0.005), or HbSS versus HbAS (Mann Whitney test, P <0.0003) **(table 1)**. Patients with abnormal HbSS phenotypes had lower hematological parameters than those with HbAS and HbSC forms. Anemia (Hb < 11 g/dl blood) was found in patients with sickle cell phenotypes AS, SC and SS, and was more severe for SS phenotypes (6.51 ± 2.24 g/L) compared to AA (11.28 ± 2.10 g/L) (Mann Whitney test, p < 0.001).

**Table 1:**
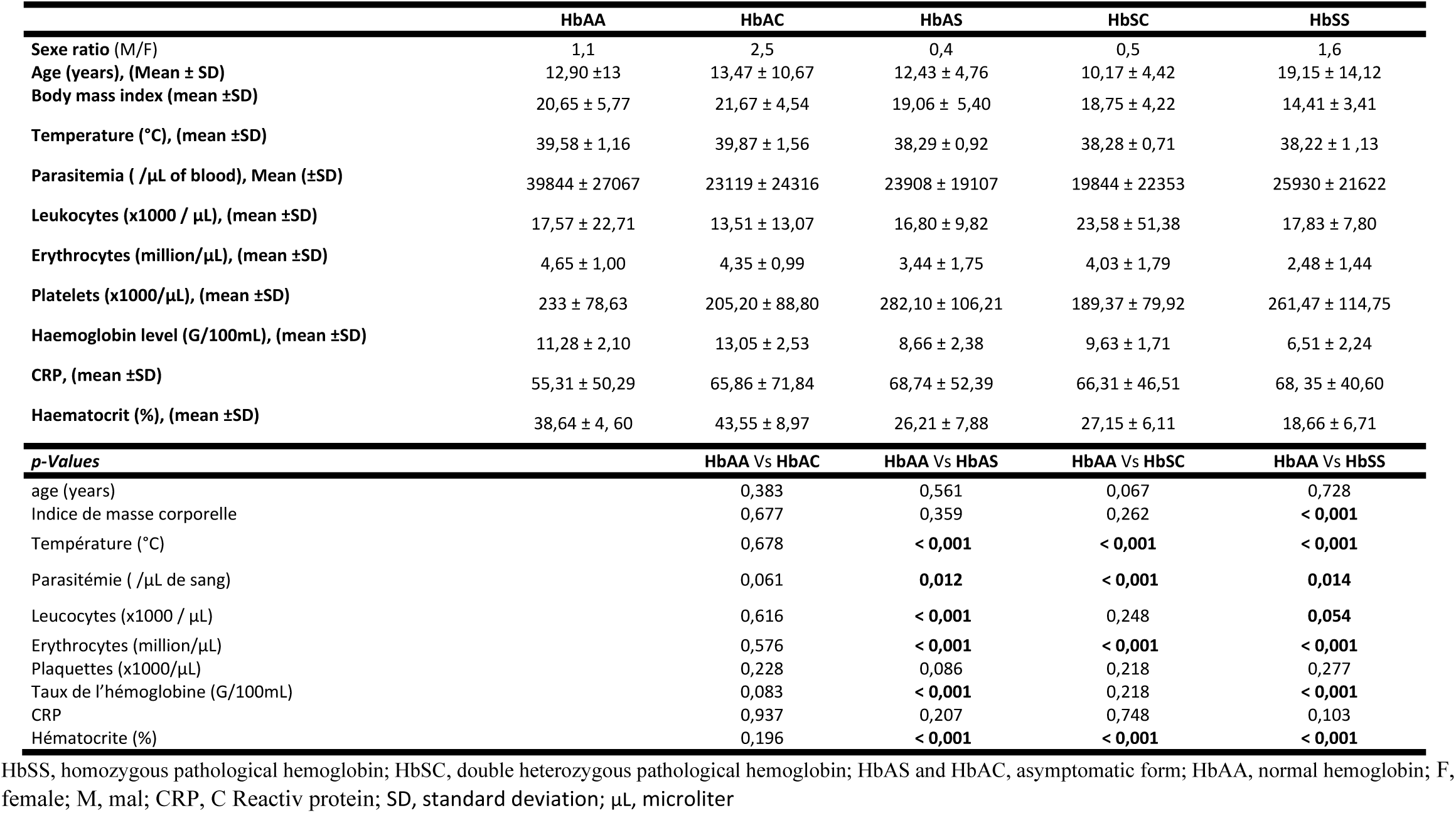
Basic parameters of the study population

|  | HbAA | HbAC | HbAS | HbSC | HbSS |
| --- | --- | --- | --- | --- | --- |
| Sexe ratio (M/F) | 1,1 | 2,5 | 0,4 | 0,5 | 1,6 |
| Age (years), (Mean ± SD) | 12,90 ± 13 | 13,47 ± 10,67 | 12,43 ± 4,76 | 10,17 ± 4,42 | 19,15 ± 14,12 |
| Body mass index (mean ±SD) | 20,65 ± 5,77 | 21,67 ± 4,54 | 19,06 ± 5,40 | 18,75 ± 4,22 | 14,41 ± 3,41 |
| Temperature (°C), (mean ±SD) | 39,58 ± 1,16 | 39,87 ± 1,56 | 38,29 ± 0,92 | 38,28 ± 0,71 | 38,22 ± 1,13 |
| Parasitemia ( /μL of blood), Mean (±SD) | 39844 ± 27067 | 23119 ± 24316 | 23908 ± 19107 | 19844 ± 22353 | 25930 ± 21622 |
| Leukocytes (x1000 / μL), (mean ±SD) | 17,57 ± 22,71 | 13,51 ± 13,07 | 16,80 ± 9,82 | 23,58 ± 51,38 | 17,83 ± 7,80 |
| Erythrocytes (million/μL), (mean ±SD) | 4,65 ± 1,00 | 4,35 ± 0,99 | 3,44 ± 1,75 | 4,03 ± 1,79 | 2,48 ± 1,44 |
| Platelets (x1000/μL), (mean ±SD) | 233 ± 78,63 | 205,20 ± 88,80 | 282,10 ± 106,21 | 189,37 ± 79,92 | 261,47 ± 114,75 |
| Haemoglobin level (G/100mL), (mean ±SD) | 11,28 ± 2,10 | 13,05 ± 2,53 | 8,66 ± 2,38 | 9,63 ± 1,71 | 6,51 ± 2,24 |
| CRP, (mean ±SD) | 55,31 ± 50,29 | 65,86 ± 71,84 | 68,74 ± 52,39 | 66,31 ± 46,51 | 68,35 ± 40,60 |
| Haematocrit (%), (mean ±SD) | 38,64 ± 4,60 | 43,55 ± 8,97 | 26,21 ± 7,88 | 27,15 ± 6,11 | 18,66 ± 6,71 |
| **p-Values** |  | **HbAA Vs HbAC** | **HbAA Vs HbAS** | **HbAA Vs HbSC** | **HbAA Vs HbSS** |
| age (years) |  | 0,383 | 0,561 | 0,067 | 0,728 |
| Indice de masse corporelle |  | 0,677 | 0,359 | 0,262 | **< 0,001** |
| Température (°C) |  | 0,678 | **< 0,001** | **< 0,001** | **< 0,001** |
| Parasitémie ( /μL de sang) | | 0,061 | **0,012** | **< 0,001** | **0,014** |
| Leucocytes (x1000 / μL) | | 0,616 | **< 0,001** | 0,248 | **0,054** |
| Erythrocytes (million/μL) | | 0,576 | **< 0,001** | **< 0,001** | **< 0,001** |
| Plaquettes (x1000/μL) | | 0,228 | 0,086 | 0,218 | 0,277 |
| Taux de l'hémoglobine (G/100mL) |  | 0,083 | **< 0,001** | 0,218 | **< 0,001** |
| CRP |  | 0,937 | 0,207 | 0,748 | 0,103 |
| Hématocrite (%) |  | 0,196 | **< 0,001** | **< 0,001** | **< 0,001** |

HbSS, homozygous pathological hemoglobin; HbSC, double heterozygous pathological hemoglobin; HbAS and HbAC, asymptomatic form; HbAA, normal hemoglobin; F, female; M, mal; CRP, C Reactiv protein; SD, standard deviation; $\mu$ L, microliter

### Ring stage essay (RSA) and maturation test for sickle cell phenotypes

RSA and maturation tests were conducted in parallel for 134 and 116 patients respectively. Only 79.85 % (107/134) and 95.69 % (111/116) of RSA and maturation test were respectively successful (**Figure 1**). Indeed, a low rate of parasitic growth occurred more often with AS red cell (**Figure 2A**). RSA values for all collected clinical isolates varied from 0 to 33.75 %. According to a threshold of 1%, 81.31% (87/107) of the clinical isolates were sensitive (< 1%), and 18.69% (20/107) resistant (> 1%) to DHA (**Table 2**).

**Figure 2:**
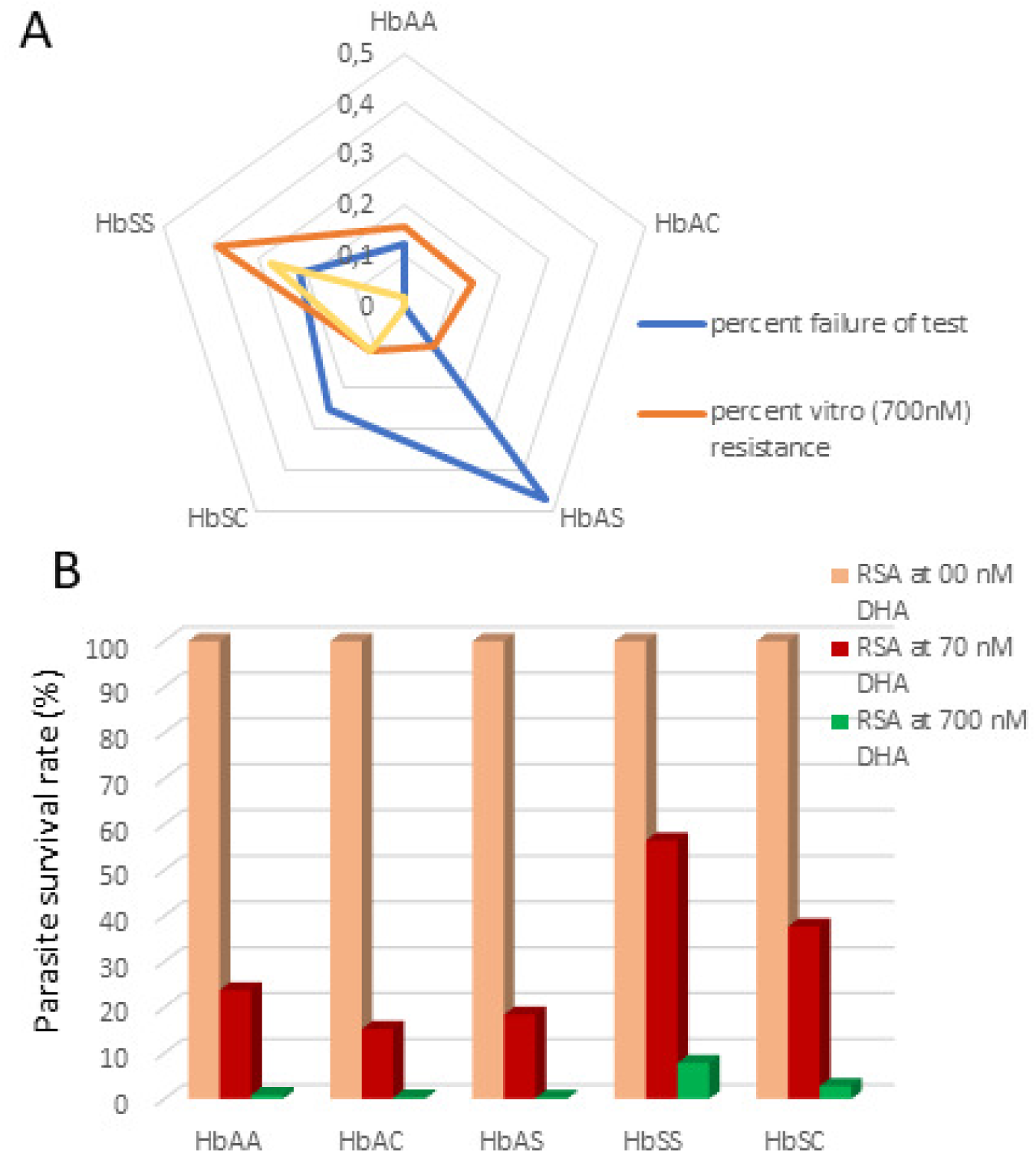
Results of the RSA test. A/ percent of test failure according to the type of hemoglobin B/ Mean parasite survival rate according to the type of hemoglobin

**Table 2:**
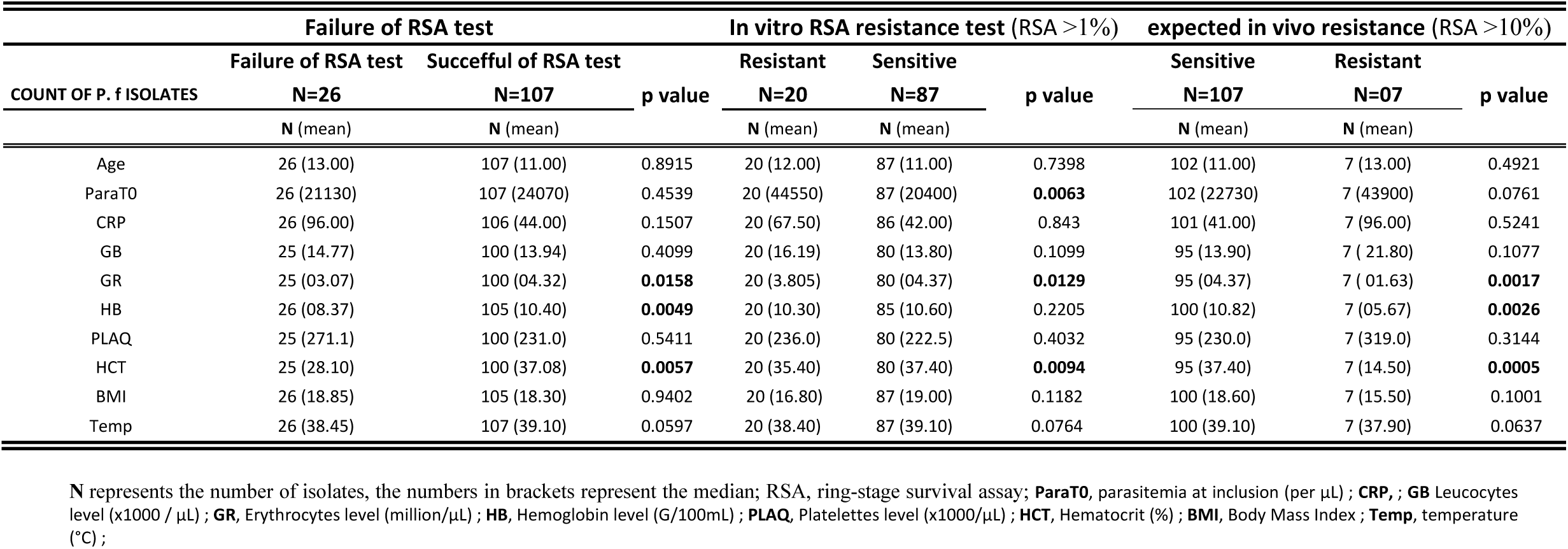
Relationship between RSA test results and patients’ clinical-biological parameters.

For the 20 resistant clinical isolates to DHA, the RSA values varied from 1.28 to 33.75 %. Resistance was found for all sickle-cell phenotypes but 39 % of the clinical isolates with a limit sensitivity at 700 nM DHA were for SS group **(Figure 2A)**. Indeed, 28% of clinical isolates from SS phenotype had survival rates ranging from 14.68% to 33.75%. (**Figure 2A**). These clinical isolates had higher ring stage parasitaemia at inclusion (70% to 75%). Parasites infesting HbSS hemoglobin are thus more resistant in culture than parasites infesting HbAS, HbAC, HbSC and HbAA **(figure 2B)**.

To confirm adequate culture conditions, the RSA tests were performed with *P falciparum* K1 strain (artemisinin sensitive strain) and IPC 3445 (Cambodian strain resistant to artemisinin) as negative and positive control respectively. K1 parasite had 0 % survival whereas IPC 3445 had 23.14%. Three clinical isolates from SS patients exhibited higher survival rates than the IPC 3445 strain **(figure 3A)**.

**Figure 3:**
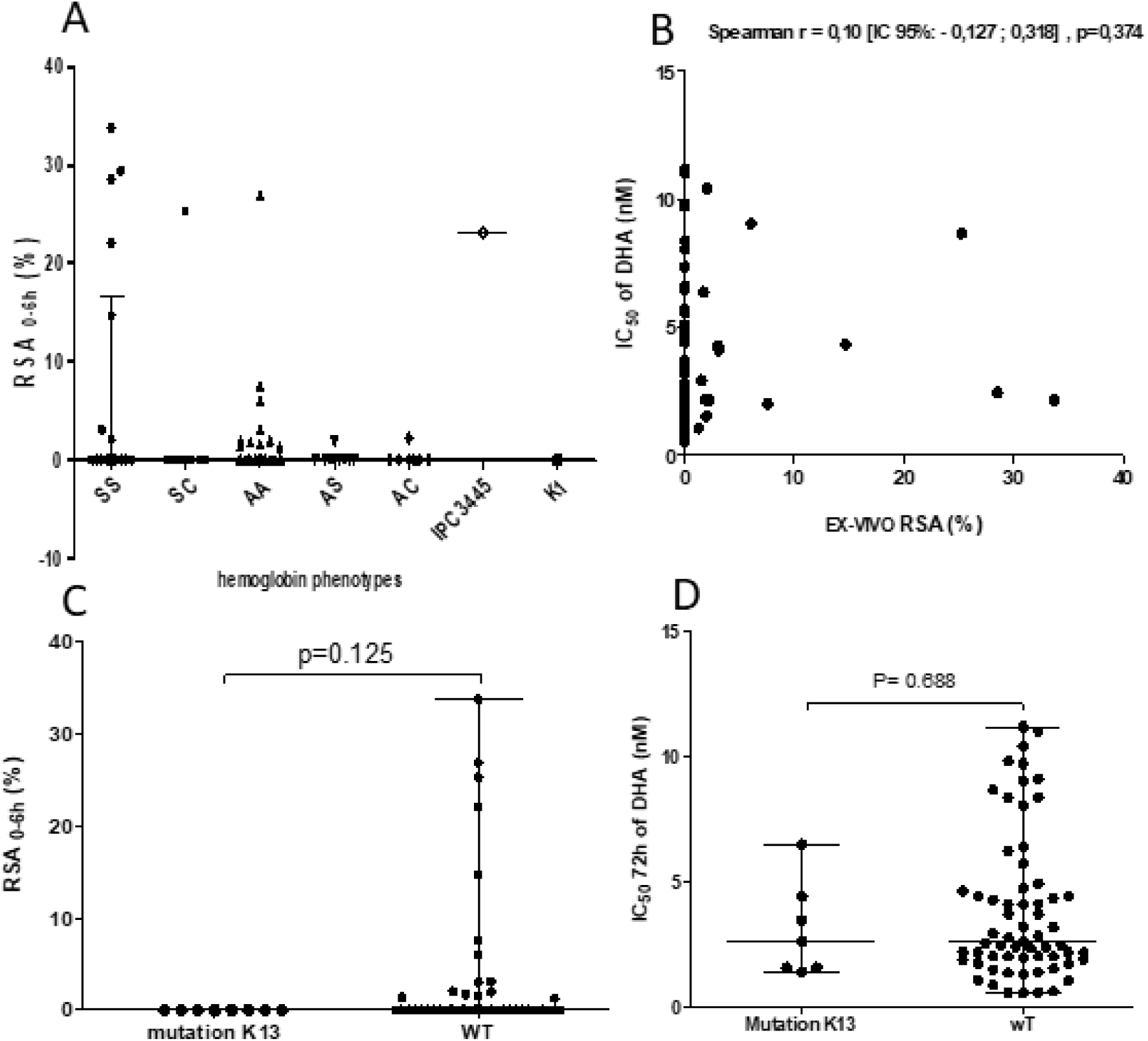
In vitro Modulation of parasites drug sensitivity. A/ Distribution of *Plasmodium falciparum* survival rates by ex-vivo RSA according to the forms of sickle cell disease. Survival rate was calculated as follows: (parasitemia at 700 nM dihydroartemisinin exposed / parasitemia at 00 nM control) × 100. RSA, ring-stage survival assay. Geometric means with 95% confidence intervals of the survival rates are shown B/ speaman’s correlation between Ex-vivo RSA (%) and IC50 values for dihydroartemisinin C/ Correlation of parasite survival rates (Ex-vivo RSA) and K13 polymorphisms. RSA values between parasites with mutations in the kelch propeller domain (> 440 aa) and parasites without kelch mutations. D/ Comparaison of CI50 values for DHA between the group of K13 wild-type parasite and the group within the propeller domain (> 440 aa)

In the same way, 85.6% (95/111) of the *in vitro* standard maturation test were successful, with Inhibition Concentration 50% (IC_50_) ranging between 0.53 and 11.18 nM (geometric mean, 2.71 nM, IC [2,32 - 3,17], range [0,53 - 11,18]) for a threshold of resistance to DHA at 10 nM.. With the RSA, most of the clinical isolates not responding to DHA were from SS patients. (**Figure 3A)**. However, when comparing both tests with the same isolates (**Figure 3B**), some isolates with a DHA-sensitive phenotype in the RSA expressed a resistant phenotype in maturation test and vice versa. As already published, the pairwise comparison of the two tests was not significant (n=81, Spearman r = 0.10 [95% CI: -0.127; 0.318], p=0.374).

### Relations between RSA test results and patients clinical-biological parameters

In order to take into account confounding variables, quantitative parameters of hosts (age, body mass index, CRP, hemoglobin level, hematocrit) were compared firstly with parasite growth rates, and secondly between groups with *in vitro* susceptibility or resistance (below and above 1%) and finally between *in vivo* sensitivity and resistance (10 % survival rate as a threshold) (**Table 2**).

Red blood cell count, hemoglobin level and hematocrit level at inclusion were significantly lower in clinical isolates with insufficient parasite maturation, i.e. failure of the test, compared to clinical isolates with growth rate > 1% (Mann Whitney test, P = 0.016, P = 0.005, and P = 0.006 respectively). **(table 2)**.

In the same way, patients with DHA-resistant isolates had a lower red blood cell count and hematocrit level compared to DHA-sensitive isolates. DHA-resistant isolates had also a significantly higher parasitemia at inclusion than DHA-sensitive isolates (44778 ± 22347 Vs 30795 ± 26671; Mann Whitney test, P= 0.006). This difference was also found for patients with HbSS (parasitemia 48550 ± 21145 Vs 16486 ± 14401; Mann Whitney test, P = 0.0021 for resistant versus sensitive isolates). For SS patients, DHA-resistant isolates had also a significantly higher parasite growth rate compared to sensitive clinical isolates (5.04 ± 4.92% Vs 1.57 ± 0.44%; Mann Whitney test, P= 0.0002). **(table 2)**. Overall, parasite development in HbSS red blood cell was adequate, but the response to DHA was lower. For the *in vivo* resistance isolates (survival rate above 10%), low red blood cell count, hemoglobin level and hematocrit at inclusion were observed only for sickle cell phenotype SS (Mann Whitney test, P = 0.0017, P = 0.0026, P = 0.0005 respectively) **(table 2)**.

### Survival rate and point mutations in the PfKelch 13 propeller gene

Genomic DNA was obtained from 134 isolates and an 849 bp PCR fragment corresponding to the Kelch13 Propeller region was amplified and sequenced. Polymorphism analysis was possible for 74% (99/134) of these sequences. Overall, 16 SNPs (Single Nucleotide Polymorphisms) were detected for only 7% of the sequences (i.e., 7/99). Among these SNPs, 94.44% were non-synonymous mutations (15/16). The synonymous mutation (G287G) was located before the propeller region of the K13 gene (< 442 amino acids), while the 15 non-synonyms were located in this propeller region. Mutations were all different (**Table 3)**. No key mutations already identified in the Kelch 13 propeller domain by other authors (such as C580Y, R539T, Y493H, P574L, I543T, F446I, R561H, A675V) and associated with a delay in parasite clearance was found. Only one mutation was found for patients with HbSS or HbAC phenotypes. No mutation was associated with a decreased drug susceptibility both for RSA and standard maturation test. No difference in IC50 values for DHA was found between isolates with and without mutations (Mann Whitney U test, P = 0.688) (**Figure 3A-B**).

**Table 3:**
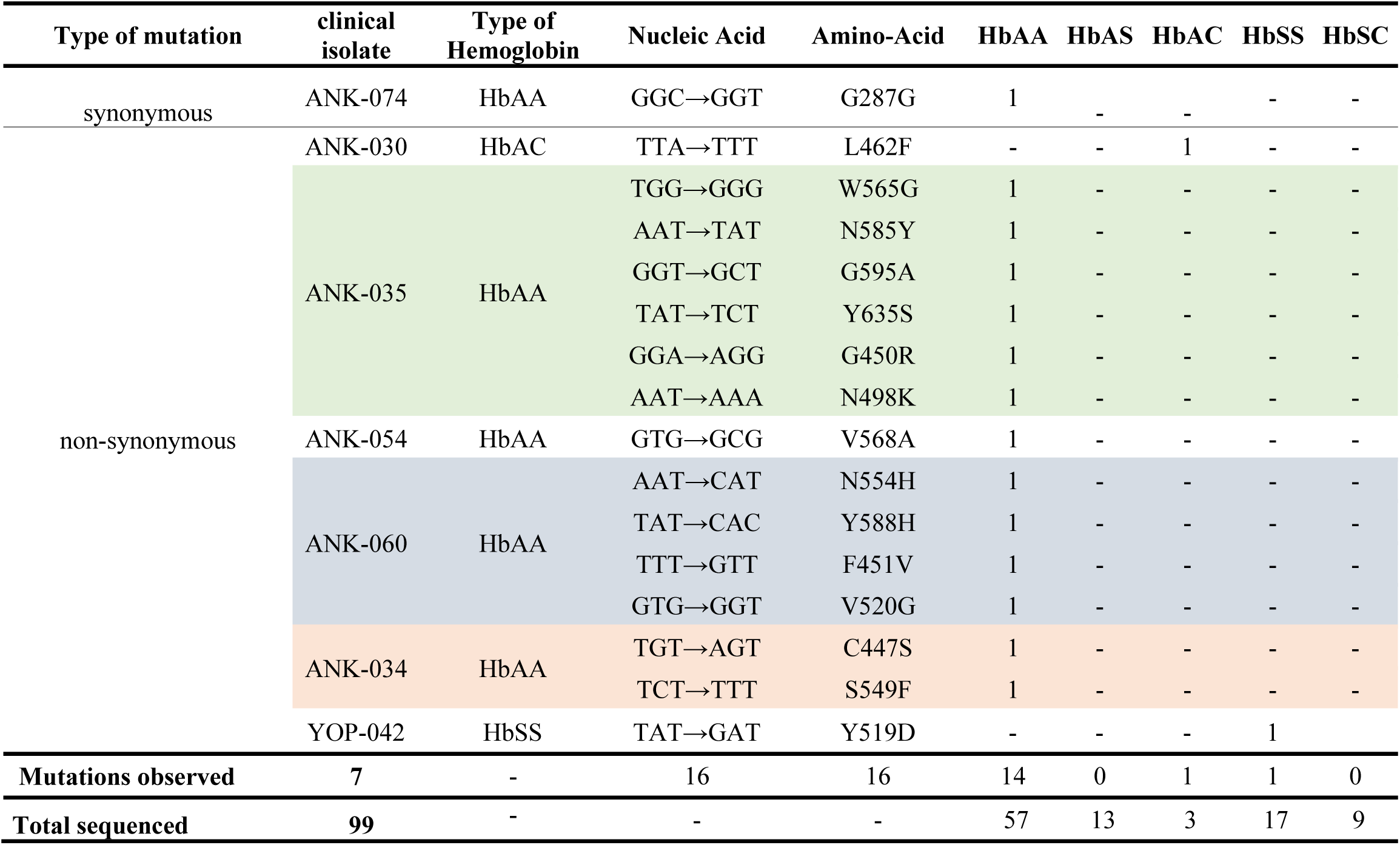
mutations identified in klech13 propeller of plasmodium falciparum in sickle cell disease patients, Côte d’lvoire.

## DISCUSSION

*In vitro* tests and gene polymorphism analysis with in *vivo* clinical studies, can serve as predictive markers for epidemiological surveillance of artemisinin resistance (31, 48, 49). However, very few studies address sensitivity of isolates contaminating patients with abnormal hemoglobin. This question is of importance, as in Côte d’Ivoire almost 20% of the population is carrying at least a genetic trait. Homozygote patients are at risk of severe occlusive crisis when infected with malaria. In the same time, cytosolic content of the abnormal red blood cells (64–66) can provide a specific biochemical environment susceptible to select or promote selection of ACT resistant isolates.

The participants included in the study were divided into four groups according to their sickle cell phenotypes (HbSS, HbSC, HbAS, HbAC and HbAA), and no significant difference in mean age was found, underlining the efficacy of the clinical management of children with SCD at the YUH (13, 50–54). As already reported, parasitaemia and haemoglobin levels at inclusion were lower in sickle cell patients with major forms than in patients with normal phenotype (55–58). This low parasite density could be an element explaining a protective effect against severe malaria. It could be due i) to dehydration of red blood cells which could inhibit the invasion and growth of *P. falciparum* parasites (57, 59, 60); or ii) to inhibition of osmotic shock in SS phenotype erythrocytes (61) resulting in reduced merozoite release (62, 63).

Due to their particular intra-erythrocyte microenvironment (64–66) attention must be paid to confirm viability of the parasites during the *in vitro* culture. It’s why a lower dose of drug i.e., 70 nM was introduced as control.

Overall the survival rates obtained during RSA test, showed higher values for isolates from patients with SS phenotype than others. These data suggest that these isolates could be resistant to DHA *in vitro* (survival rate >1%), and *in vivo* (survival rate >10%). These values could be due to a higher density of ring stages (70-75%) before culture which are known to enter quiescence in the presence of DHA (25, 27–29). However, a recent work in Abidjan underlined as well a higher complexity of the parasite isolates in patients with SCD or trait compared with control ones (37). This can point out selection of a specific set of parasites entering abnormal red cells. To test this hypothesis, sequencing of the full genome of these parasites is in process. In the same line two other hypothesis could explain a decrease in ACT sensitivity of these parasites. Resistance could be the result of an adaptation to novel micro-environmental or biochemical conditions (18, 19, 66, 74) with a different transcriptome regulation and activation of specific genes (30). Gene expression analysis studies should also be conducted. Inactivation of DHA *in situ* by the local biochemical conditions could also be postulated. Nevertheless, mutations in the K13 gene (in particular the WHO-validated C580Y, R539T, Y493H, P574L, I543T, F446I, R561H, A675V, N458Y (67)) are not associated with decrease in sensitivity. This absence of link between K13 mutations and DHA sensitivity was already described elsewhere in Africa, as in Cameroon (34) and Uganda (35). and even in Cambodia (68).

To the best of our knowledge, this work demonstrates for the first time *in vitro* resistant isolates in SS patients not related to polymorphism in the propeller region of the PfKelch 13 gene.

This resistance could be explained by activation of alternative metabolic pathways as unfolded protein pathways seem up-regulated to attenuate artemisinin-induced protein damage (30). In the same line several other genes could be involved in this resistant phenotype as falcipain 2a (FP2a) a cysteine protease and haemoglobinase. Mutations in this enzyme (FP2a) reduce enzymatic activity and haemoglobin digestion, and increase the survival rate in the ring stage of P. falciparum (69, 70). Mutations outside the K13 Propeller gene could also induce compensation effect as already reported with the Pfcrt gene in French Guiana (71-73).

However, this study supports overall the idea that isolates from SS sickle cell patients, can express DHA resistant phenotype without Kelch 13 Propeller gene polymorphism (34, 48, 67, 79–81).

## CONCLUSION

The chemoresistance of *Plasmodium falciparum* to antimalarial drugs makes malaria control more complex. The results of this study emphasize a worrying increase in *in vitro* DHA-resistant strains, particularly in SS sickle cell patients. This study also provides evidence of no relation between Pfkech13 polymorphism and survival rate in RSA in sickle cell patients living in Abidjan. Taken together, these results highlight the need for appropriate and effective treatment in these subjects to protect them from severe attacks and to avoid the circulation of resistant strains.

## MATERIALS and METHODS

### Site of study

This prospective study was carried out from May 2017 to February 2019 at the department of Clinical Hematology at Yopougon University Hospital (YUH, Abidjan) and at the community health center of Anonkoua-kouté (ANK, Abidjan). YUH is the reference center for sickle cell disease in Côte d’Ivoire where about 9,000 patients with major sickle cell disease are followed up with free access to medical care. The community health center of Anonkoua-kouté is a secondary level health structure which receives more than 400 patients daily. Patients attending this health center can benefit from the typing of hemoglobin using acid acetate electrophoresis.

### Patients recruited and samples collection

For patients suffering from fever attending both health structures, a clinical examination was performed before a biological confirmation of malaria. A first screening was carried out by lateral flow test. Positive results were validated by examination of thick and thin blood Giemsa stained smears at x1000 with light microscopy. For positive sample, parasite density was calculated. After written informed consent of participants or of their legal representatives, all patients over 6 months of age with a parasite density beyond or equal to 0.1 % were included in the study. An electrophoresis of hemoglobin was performed to all the patients registered. For enrolled patients, a questionnaire was applied including demographic data, sex, age, place of residence, body temperature and clinical symptoms.

Patients with signs of severe malaria (WHO criteria) and/or requiring intensive medical care for other severe diseases, as well as those already treated with antimalarial drugs within the 30 days prior to medical consultation were not include and directly sent to physician consultation with their biological results.

In YUH only patients with already known sickle cell disease and malaria were recruited. Whereas in Anonkoua-koute health center all the patients with positive thick blood smears were enrolled after informed consent notwithstanding the result of the electrophoresis.

For each patients 3 mL of peripheral venous blood was collected on EDTA tubes for culture and 2 ml of blood were collected on dry tubes for biochemical tests (CRP). Blood spots were also done with three drops (50 µl each) put on a Whatmann 3MM® filter paper and dried at room temperature for 4 hours. Samples were kept at 4°C in an ice chest cooler and sent to Institut Pasteur of Côte d’Ivoire malaria unit in less than four hours.

This study protocol was approved by the National Ethics and Research Committee of the republic of Côte d’Ivoire.

### Hemoglobin status

Patients enrolled at YUH were already aware of their sickle cell status and were all carriers of major forms (SS, SC and CC). These patients were routinely treated and followed up by the reference center. Nevertheless, they genetic status was validated by FRET method (83). At Anonkoua-Kouté Health Center, screening for sickle cell disease diagnosis was done by electrophoretic using an SAIO Electrophoresis instrument.

### *in vitro* drug sensitivity test

*In* vitro tests were performed using RPMI-1640 (Eurobio 479604, 500ml) medium supplemented with 5% Albumax II, 1% L-glutamine, 2% D-glucose, 0.05% hypoxanthine, 2.5% HEPES (Eurobio 251010) buffer and 0.5 % gentamicin (Eurobio 524221). Serum and buffy coat were removed from the whole blood obtained from patients and red blood cells were washed three times in RPMI-1640 medium (centrifugation at 3000 rpm for 10 minutes) prior to cultivation. Samples were seeded in culture less than 5 hours after blood collection. The cultures were conducted in a modular incubator chamber saturated with 5% O_2_, 5% CO_2_ and 90% N_2_ in a humidified atmosphere.

### Ring Stage Assay

The test was conducted according to Witkowski et al. (84) with minor modifications. To confirm viability of clinical isolates, two concentrations of di-hydro-artemisinin (DHA) were used for each isolate, i.e. 700 nM and 70 nM. Dimethylsulfoxide (DMSO) at 0.1% was used as negative control.

The rest of the procedure did not change. Briefly the modular incubator chamber was placed in an incubator at 37°C for six (6) hours. After six (6) hours of exposure to DHA, the red blood cells were washed three times with a preheated RPMI 1640 medium and suspended in a new complete medium. Cultures were incubated under the same conditions for sixty-six (66) hours. Light microscopy Giemsa stained of thin blood smears were prepared and examined. The number of infected red blood cells containing viable parasites in a total of at least 10,000 red blood cells was counted. Viable parasites were counted to determine the survival rate. The test was considered as interpretable when at 72h, the proportion of viable parasites in culture wells not exposed to DHA is higher than the initial parasitaemia at T0 (84). Parasite isolates demonstrating > 1% survival in the RSA were considered to display reduced susceptibility to ART (85).

### Maturation inhibition assay

The *in vitro P. falciparum* maturation test was conducted as developed by Trager and Jensen (1976), standardized by Lebras et al and modified for fluorescent detection by Smilkstein et al., 2004 and Basco, 2007. Parasitized red blood cells were seeded in complete medium at hematocrit of 2%. In case of parasitaemia greater than 0.3%, type O positive washed healthy human erythrocytes were added to adjust parasitaemia at 0.3%. Di-hydro-artemisinin was added in duplicate in 96-well microtiter plate at concentrations ranging from 35.16 nM to 0.55 nM. As previously, incubation was conducted at 37°C for 72 hours in a modular incubator chamber saturated with 5 % O_2_, 5% CO_2_ and 90 % N_2_ in a humidified atmosphere. After incubation, cultures were frozen for 24 hours to stop parasite growth and to lyse red cells. Cultures were thawed and parasite growth was assessed by SYBR Green I incorporation method using a spectrofluorometer (DELL, FLx800, biotek) according to Smilkstein et al. (2004), Basco et al (2007) and Le Nagard et al. (2011). Drug concentrations inhibiting 50% of the parasite growth (IC50) were determined using IVART (In Vitro Analysis and Reporting Tool) software from WWARN’s (86). Validation of each test was also assessed with IVART. Resistance thresholds of DHA (10 nM) was defined according to IVART.

### K13-propeller gene sequencing

Parasitic DNA was extracted using a Qiagen kit according to the manufacturer’s instructions. The fragment 1279-2127 of the coding sequence of *k13-propeller* gene of *P*.*falciparum* was amplified by nested PCR according to Ariey et *al* (31). PCR products were sequenced according to Sanger method (Great Britain) by Genewiz compagny. Sequences aligned by Seaview 5 were analyzed using BioEdit software version 7.0.9.1 and compared to *k13-propeller* sequence (XM_001350122.1).

### Statistical analysis

Statistical tests were performed using GraphPad Prism 7.0 version (GraphPad prism software Inc., San Diego, CA, USA), and Statistica v9. The Shapiro-wilk test was used to verify data normality. Medians with interquartile deviations were used for data that do not follow a normal distribution. Mann-Whitney U-test, Kruskal Wallis test and median test were used to compare groups. Correlations were determined using the Spearman test or Kendall Tau test. Comparisons were considered statistically significant when p ≤ 0.05. <ack>

## Acknowledgement

This program and AG were supported by a grant from the Rotary Foundation. We thank all the patients who agreed to participate, and the staff of YUP and Anoukouakouté for their help.

